# Transcription Factor–Mediated Reprogramming of Cancer-Associated Fibroblasts Reveals Targetable Vulnerabilities in Solid Tumors

**DOI:** 10.64898/2026.04.15.718753

**Authors:** Nan Sook Lee, Priyankan Datta, Yixuan Huang, Bella Raykowski, Xi Yu, Tianze Guo, Peixiang He, Sreejesh Moolayadukkam, Ben Yi Tew, Shigang Xiong, Chi Woo Yoon, Peter Yingxiao Wang, Christopher DeRenzo, Jacek Pinski, Ishwar K. Puri

## Abstract

Cancer-associated fibroblasts (CAFs) contribute to immune exclusion and therapy resistance in solid tumors, limiting the efficacy of chimeric antigen receptor (CAR) T cell and immune cell therapy. To overcome this, we developed a transcription factor (TF)-based strategy to reprogram prostate-derived CAFs (pCAFs) into normal fibroblast-like cells (NFs). We prioritized TFs enriched in quiescent stellate cells—Vitamin D receptor (VDR), Peroxisome Proliferator-Activated Receptor gamma (PPARγ), and p53—and selected VDR for proof-of-concept studies. Lentiviral VDR expression in pCAFs produced VDR-reprogrammed NFs (VDR-rpNFs) with reduced CAF markers, increased ATP, and suppressed TGF-β and IL6, indicating phenotypic and metabolic reversion. In both *in vitro* 3D co-cultures and *in vivo*, VDR-rpNFs disrupted tumor architecture, enhanced CAR T cell infiltration, and reduced necrosis. PPARγ- and p53-rpNFs showed similar reprogramming effects. These results suggest TF-guided fibroblast reprogramming as a viable strategy to remodel the tumor microenvironment and improve CAR T cell efficacy in solid tumors.

## INTRODUCTION

Chimeric antigen receptor (CAR) T cell therapy and other immune-based modalities have transformed the treatment of hematologic malignancies, yet their efficacy in solid tumors remains limited^1–3^. A central obstacle is the tumor microenvironment (TME), which imposes physical, metabolic, and immunological constraints that restrict immune cell infiltration and therapeutic access. Among stromal constituents, cancer-associated fibroblasts (CAFs) are particularly influential: they orchestrate desmoplastic remodeling, secrete immunosuppressive cytokines, and reinforce extracellular matrix (ECM) stiffness, thereby promoting immune exclusion and resistance to therapy^4–8^. CAFs thus function not merely as structural elements but as active drivers of therapeutic failure, making them compelling targets for intervention.

CAFs arise from diverse origins, including normal resident fibroblasts and quiescent stellate cells ^4,9,10^. In the liver, stellate cells—representative quiescent fibroblasts (qNFs)—undergo transcription factor (TF)-driven activation, transitioning into activated fibroblasts (aNFs) and myofibroblasts (myoFs)^11^. This process underlies fibrosis, a pathological wound-healing response characterized by excessive ECM deposition and stromal scarring^4,12^. Liver injury-induced fibrosis provides a conceptual framework for understanding fibroblast plasticity and its role in fibrotic pathology across tissues^11,13^.

Fibrosis exemplifies how impaired wound healing generates physical barriers that impede therapeutic access. Importantly, fibroblast activation is reversible: restoring cells to a quiescent state can decompress the stroma and improve drug and immune cell penetration^14–16^. Notably, the vitamin D receptor (VDR) functions as a master TF in pancreatic stellate cells (PSCs), reinstating quiescence and remodeling the stroma, thereby offering a molecular strategy for tumor microenvironment reprogramming^17–20^. Chronic PSC activation in cancer drives pathological matrix secretion, but targeted TF modulation suggests this state may be reversible^17,21–22^.

Despite these advances, the TF networks governing stromal reprogramming in solid tumors remain incompletely defined. Here, we propose leveraging a set of TFs enriched in anti-inflammatory, quiescent fibroblasts—VDR^17–19^, PPARγ^23–25^ and p53^26–27^—to redirect CAFs into a tumor-suppressive phenotype. Rather than ablating CAFs, our approach seeks to restore their homeostatic function, decompress the tumor microenvironment, and enhance immune cell access^4,7,28–29^. This strategy has the potential to markedly improvè the efficacy of CAR-T and related immune cell therapies in solid tumors, which account for ∼90% of cancer cases worldwide^1,2^. Beyond oncology, such stromal reprogramming may inform a conceptual framework for next-generation cancer vaccines, integrating microenvironmental cues to optimize immune priming and durability^17,30^ and extend to other fibrosis-related diseases^11,14,31^.

## RESULTS

### Engineering pCAFs with Quiescence-Associated Transcription Factors

To explore the feasibility of reprogramming human prostate cancer-associated fibroblasts (pCAFs) toward a normal fibroblast (NF)-like state, we ectopically expressed transcription factors (TFs) enriched in quiescent stellate cells. As an initial proof-of-concept, we focused on the Vitamin D Receptor (VDR), given its well-established role in suppressing stellate cell activation (Fig. 1).

**Fig. 1.**
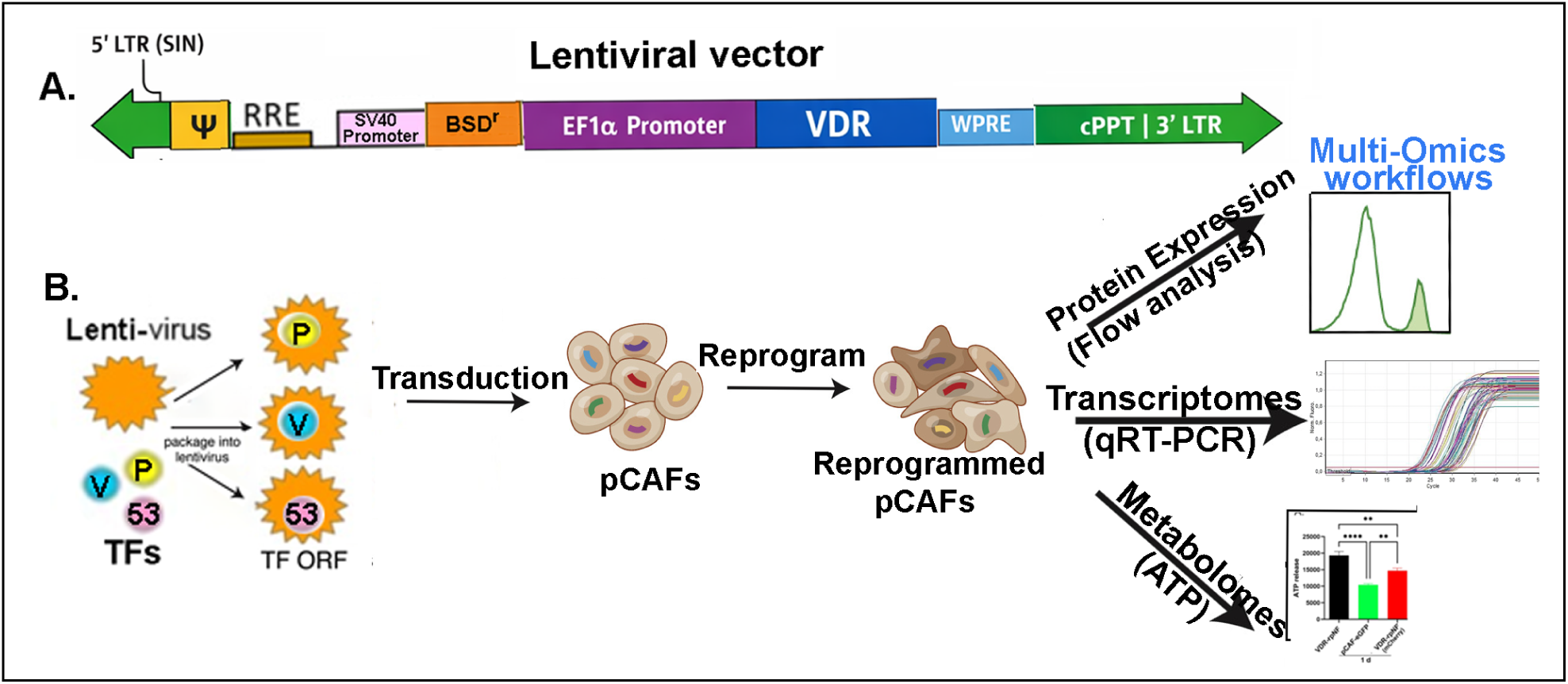
Building transcription factor modules for directed reprogramming. **A.** Lentiviral vectors used for direct reprogramming of pCAFs. The constructs incorporated SIN-modified 5′ and 3′ LTRs for safe genomic integration, a Ψ packaging signal, and an EF1α promoter driving VDR (or PPARγ or p53) expression. A BSDʳ cassette enabled Blasticidin selection, or mCherry for FACS selection, while the cPPT together with the 3′ LTR facilitated efficient nuclear import and proper transcriptional termination. **B.** Validation of individual transcription factors for reprogramming pCAFs into quiescent fibroblasts. TFs: P; PPARγ, V; VDR, 53; p53. Transduced and reprogrammed pCAFs were characterized using multi-omics workflows: Expression of pCAF-associated and normal fibroblast markers was quantified at the mRNA level by qRT-PCR and at the protein level by FACS in parental pCAFs and VDR-reprogrammed fibroblasts (rpNFs). Metabolic reprogramming was assessed by measuring ATP production in VDR-rpNFs using bioluminescent assays.

To generate stable VDR-expressing human pCAFs, we employed complementary antibiotic- and fluorescence-based enrichment strategies. Human pCAFs (hTERT PF179T; ATCC) were transduced with high-efficiency lentiviral vectors encoding VDR together with either a Blasticidin resistance cassette or fluorescent reporters (mCherry or eGFP) (Fig. 1A). Lentiviruses were produced in HEK293T cells under standard conditions and applied to pCAFs at high multiplicity. Following transduction, Blasticidin selection (2–3 µg/mL for five consecutive days) efficiently eliminated non-transduced parental cells, yielding a stable VDR-expressing population designated ‘pCAF/VDR-BSD^r^’ (Fig. 1B). Surviving cells were expanded for three passages to establish uniform cultures. In parallel, fluorescence-activated cell sorting (FACS) was used to isolate VDR-mCherry–positive pCAFs (‘pCAF/VDR-mCherry’), with eGFP-expressing cells serving as matched controls (‘pCAF/eGFP’). Together, these orthogonal enrichment approaches enabled the generation of robust, well-defined VDR-expressing pCAF populations for downstream phenotypic and functional analyses.

To broaden the scope of this approach, we extended the lentiviral transduction strategy to additional TFs implicated in the inhibition of stellate cell activation. Specifically, p53 and PPARγ were introduced into pCAFs, generating stable populations designated ‘pCAF/p53-BSD^r^’ and ‘pCAF/PPARγ-BSD^r^’. Together, these engineered cell lines provided a platform to systematically evaluate the capacity of distinct TFs to revert pCAFs toward a quiescent, NF-like phenotype.

### Characterization of Reprogrammed Fibroblasts

We next assessed the phenotypic and molecular consequences of TF expression using transcriptomic (qRT-PCR), proteomic (flow cytometry, western blot), and metabolic (ATP quantification) profiling (Fig. 1B). Analyses focused primarily on VDR-transduced cells (‘pCAF/VDR-BSD^r^’ and ‘pCAF/VDR-mCherry’) (Fig. 2A, B).

**Fig. 2.**
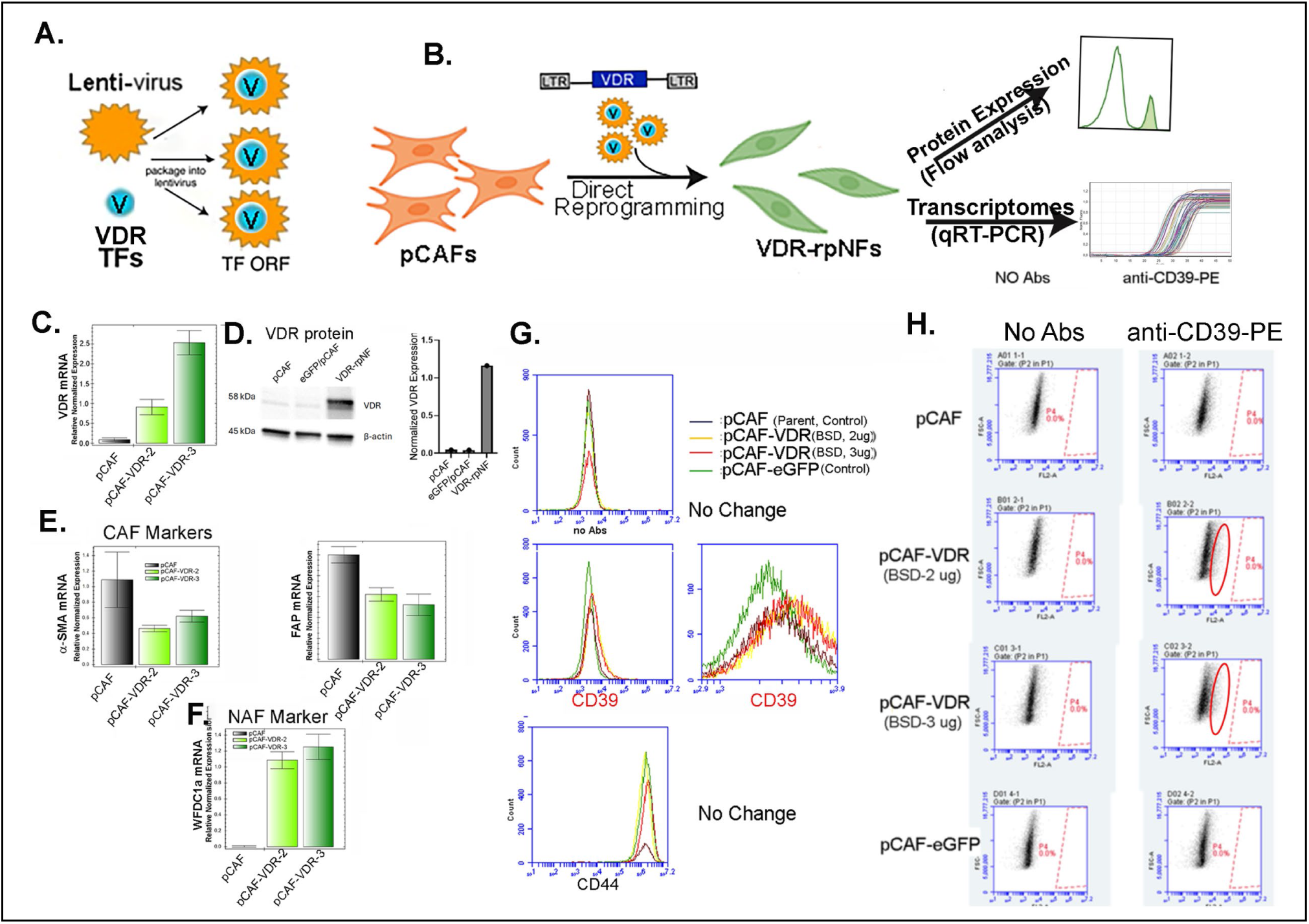
Characterization of VDR-reprogrammed pCAFs (pCAF/VDR-BSD^r^). **A, B.** Schematic of VDR transcription factor transduction into pCAFs and validation of reduced canonical CAF activation markers by qRT–PCR and FACS. **C, E, F.** qRT–PCR, **D.** Western blot, and **G, H.** FACS analyses of pCAFs transduced with VDR alone. VDR expression induces partial suppression of CAF markers and increased expression of quiescent NF-associated markers. For qRT–PCR analyses, gene expression was normalized to GAPDH and reported as relative mRNA levels in control pCAFs and VDR-rpNFs (C, E, F). Bar graphs show biological replicates plotted as mean ± s.e.m.; statistical significance was assessed by one-way ANOVA with Friedman’s test for multiple comparisons. For Western blots, a representative image from n = 3 independent experiments is shown (D). FACS data represent mean fluorescence intensity (MFI) or percentage-positive cells from biological replicates (N=3), shown as mean ± s.e.m. Statistical analyses are indicated in the panel.

pCAF/VDR-BSD^r^ cells showed strong induction of VDR mRNA and protein, whereas parental pCAFs lacked detectable expression (Fig. 2C, D). Blasticidin dose–response analysis revealed 10-fold and 25-fold increases in VDR expression at 2 µg/mL and 3 µg/mL, respectively, relative to parental cells; 3 µg/mL was used for all subsequent studies.

Relative to parental pCAFs, pCAF/VDR-BSD^r^ cells exhibited ∼2-fold reductions in α-SMA and FAP and a >10-fold increase in the NF-associated marker WFDC1 (Fig. 2E, F), indicating partial reversion toward a quiescent fibroblast state. Flo cytometry further showed a reproducible 12–14% increase in CD39, a quiescent fibroblast marker typically downregulated in CAFs ^32^ (Fig. 2G, H). These data demonstrate that VDR overexpression reduces CAF activation markers, enhances NF-associated gene expression, and partially restores quiescent fibroblast features; we therefore refer to these cells as VDR-reprogrammed normal fibroblasts (VDR-rpNFs).

To assess whether additional TFs exert similar effects, we profiled p53-rpNFs and PPARγ-rpNFs (Fig. 3A,B). Both populations showed reduced CAF activation markers, with α-SMA and FAP mRNA decreased by ∼2–10-fold and ∼6–12-fold, respectively (Fig. 3C). PPARγ-rpNFs also exhibited a ∼3-fold reduction in FSP mRNA, whereas p53-rpNFs showed minimal change. Effective transduction was confirmed by ∼10-fold increases in PPARγ and p53 mRNA (Fig. 3D). CD39 protein levels increased modestly (15–20%) in both populations (Fig. 3E), consistent with partial phenotypic reversion. These findings indicate that individual TFs can partially reprogram pCAFs toward an NF-like state.

**Fig. 3.**
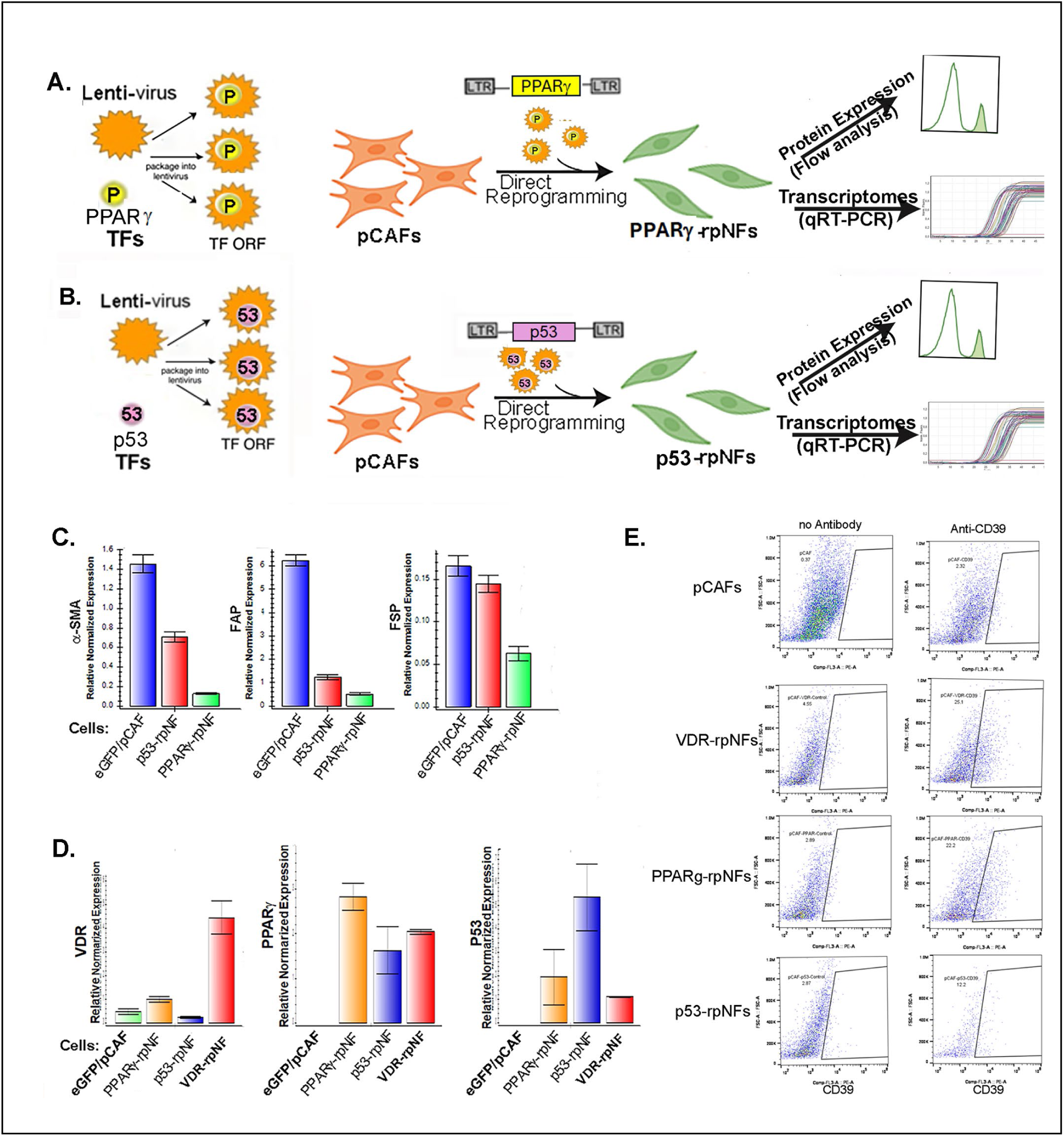
Characterization of the selected, other TFs (PPARγ and p53). **A, B.** Schematic of PPARγ and p53 transcription factor transduction into pCAFs and validation of reduced canonical CAF activation markers by qRT–PCR and FACS. **C.** Modulation of expression in CAF activation markers in PPARγ- or p53-rpNFs. **D.** Exprssion of TFs in pCAFs transduced by PPARγ and p53 by Q-RT-PCR**. E.** FACS analysis on CD39 protein expression in PPARγ-rpNFs and p53-rpNFs. For qRT–PCR analyses, gene expression was normalized to GAPDH and reported as relative mRNA levels in control eGFP/pCAFs, and PPARγ-rpNFs or p53-rpNFs (C, E, F). Bar graphs show biological replicates plotted as mean ± s.e.m.; statistical significance was assessed by one-way ANOVA with Friedman’s test for multiple comparisons. FACS data represent mean fluorescence intensity (MFI) or percentage-positive cells from biological replicates (N=3), shown as mean ± s.e.m. Statistical analyses are indicated in the panel.

### Metabolic Reprogramming of TF-Engineered Fibroblasts

Because quiescent fibroblasts rely on oxidative phosphorylation (OXPHOS), whereas activated CAFs favor aerobic glycolysis (Fig. 4A)^31,33,34^, we quantified ATP production in reprogrammed cells. VDR-rpNFs displayed markedly elevated ATP levels—85.5% ± 6.8% (Day 1), 68.3% ± 1.4% (Day 2), and 84.7% ± 5.2% (Day 4)—relative to pCAF/eGFP controls (Fig. 4B). By Day 4, PPARγ-rpNFs and p53-rpNFs similarly showed increased ATP production (83.6% ± 12.3% and 33% ± 8.9%, respectively) (Fig. 4C). These results indicate that TF-mediated reprogramming restores metabolic features characteristic of normal fibroblasts.

**Fig. 4.**
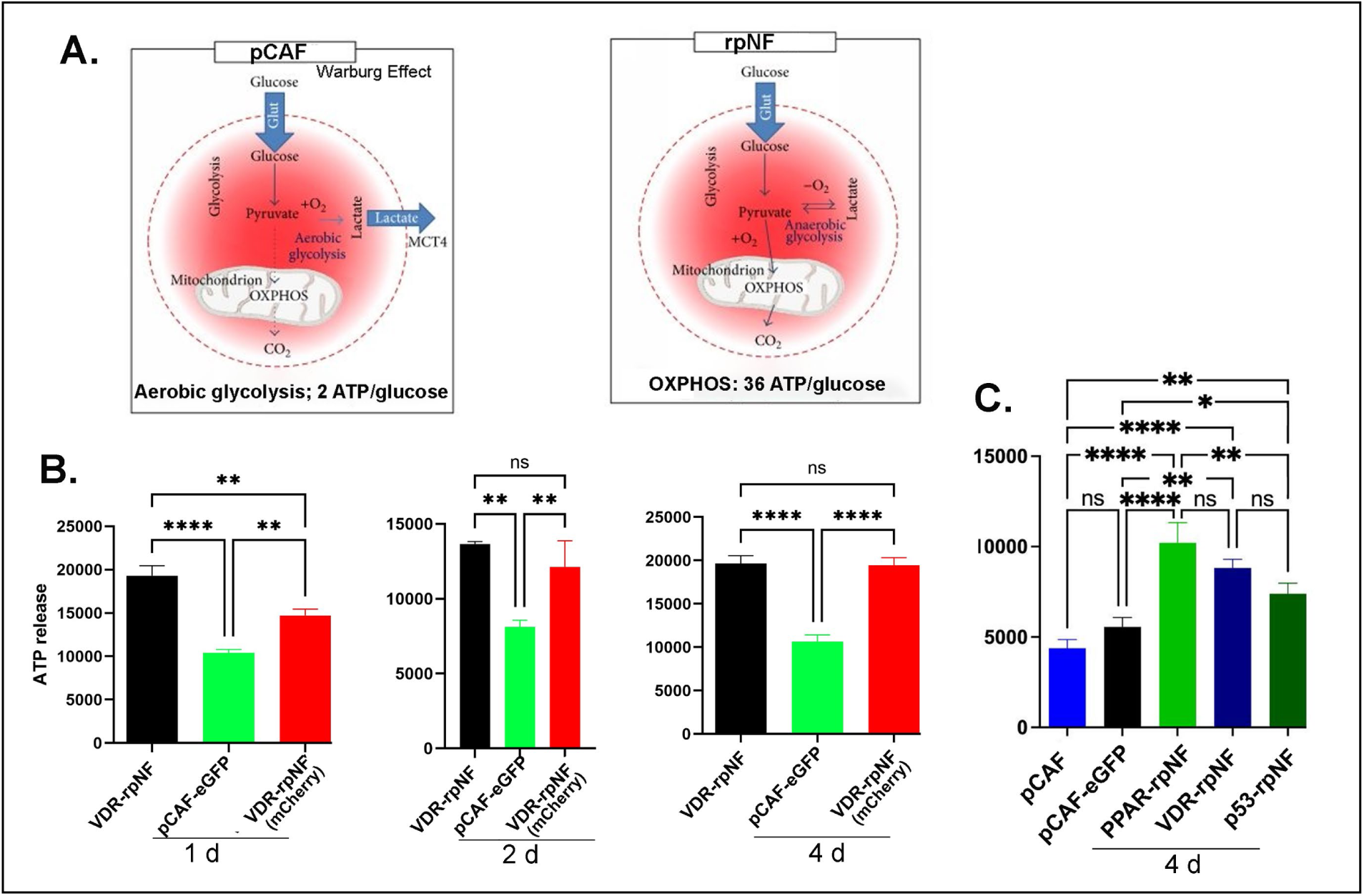
Metabolic comparison of ATP-generating pathways in pCAFs and reprogrammed normal fibroblasts (rpNFs). **A.** pCAFs show suppressed oxidative phosphorylation (OXPHOS), reduced mitochondrial ATP output, and a compensatory shift toward aerobic glycolysis, leading to elevated lactate production and export. In contrast, rpNFs display restored mitochondrial metabolism, with an active TCA cycle driving the electron transport chain (ETC) to support efficient OXPHOS and increased ATP generation. **B.** ATP quantification assays. ATP levels in VDR-rpNFs were measured at multiple time points (1-4 d) after seeding, using pCAF/eGFP cells as controls. C. ATP production across distinct rpNF lines was assessed at day 4 after seeding, with pCAF or pCAF-eGFP cells as controls. Statistical significance: **** < 0.0001; ** < 0.0033; * < 0.05.

### Suppression of Fibrotic and Inflammatory Signaling by VDR

VDR activation is known to attenuate fibroblast activation by inhibiting the TGF-β1/Smad2/3 axis^35^ (Fig. 5A). Consistent with this, VDR-rpNFs exhibited reduced expression of fibrotic genes and diminished production of inflammatory cytokines. qRT-PCR analysis revealed ∼6-fold and ∼3-fold reductions in IL-6 and TGF-β mRNA, respectively, relative to parental pCAFs (Fig. 5B). Given the roles of IL-6 and TGF-β in promoting immunosuppression and therapy resistance ^36–39^, their coordinated downregulation suggests a shift toward a less suppressive stromal phenotype that may enhance CAR T-cell infiltration and effector function.

**Fig. 5.**
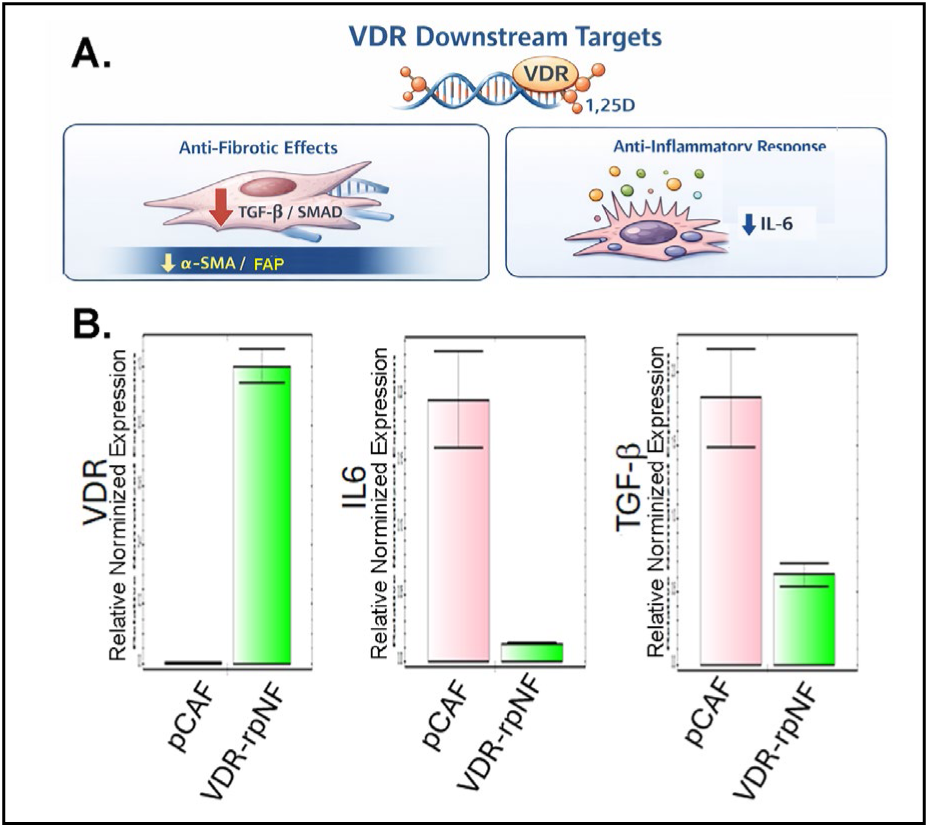
VDR downstream targets in VDR-rpNFs. **A.** Schematic representation of VDR-regulated pathways in pCAFs. VDR inhibits TGF-β/Smad2/3 signaling, a central driver of CAF activation, and suppresses inflammatory amplification loops, including reduced expression of IL-6 and other cytokines as downstream VDR targets. **B.** qRT-PCR analysis of inflammatory cytokine and TGF-β expression following CAF reprogramming. VDR-rpNFs exhibit reduced IL-6 and TGF-β mRNA levels compared with parental pCAFs. For qRT-PCR analyses, gene expression was normalized to GAPDH and reported as relative normalized mRNA levels in control pCAFs and VDR-rpNFs (C, E, F). Bar graphs show biological replicates plotted as mean ± s.e.m.

### Magnetically guided 3D bioprinting of PC-3 spheroids recapitulates hypoxic tumor-core architecture

Using a magnetically assisted 3D bioprinting platform, we generated PC-3 spheroids that reproducibly established complex tumor architecture and stromal-interaction niches. PC-3 or PC-3/GCaMP8 cells rapidly aggregated into spheroids within three hours following magnetic field–driven assembly of ∼5,000 cells per well in medium containing the paramagnetic agent Gadavist ^40–42^(Fig. 6A–C). The resulting structures displayed morphologies characteristic of hypoxic tumor cores (Fig. 6D), closely mirroring in vivo tumor microenvironments.

**Fig. 6.**
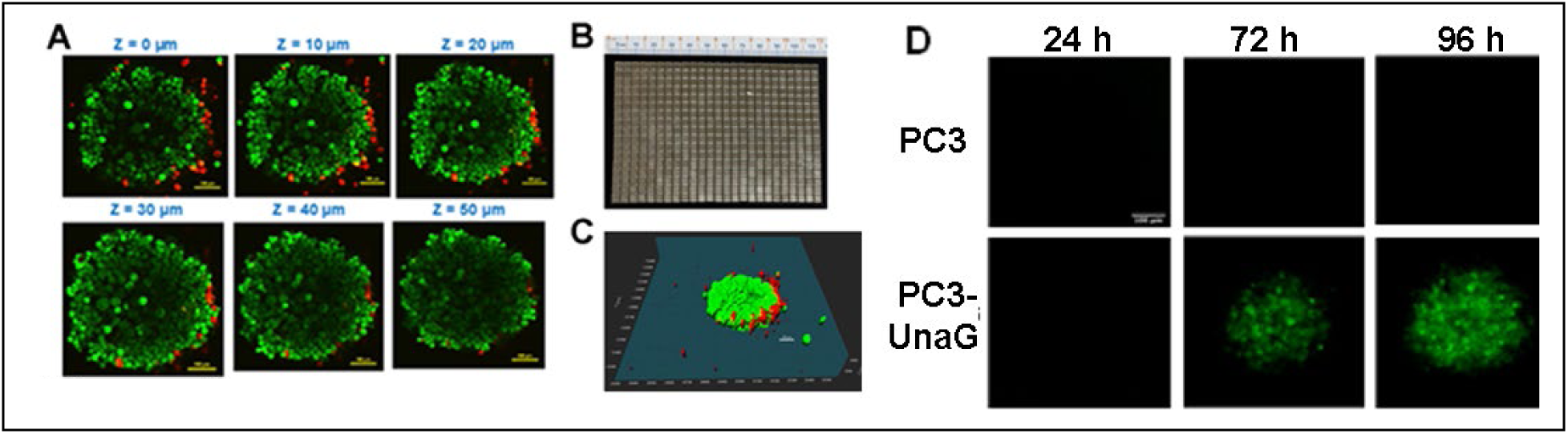
**A.** PC-3 spheroid formed using the magnetic printing technique within 3 hrs. Confocal images of the live cells after staining with Calcein AM green and the dead cells PI red at different optical planes. **B.** Picture of magnetic array containing cuboid permanent magnets. **C.** The size of a spheroid is 50-80 mm. **D.** PC-3 and PC-3-UnaG spheroids with three different time after printing exhibiting hypoxia through UnaG expression.

To verify hypoxia within the spheroid interior, we used PC-3/UnaG cells expressing hypoxia-responsive elements (HREs) fused to the oxygen-independent fluorescent reporter UnaG^43,44^. Live-cell imaging revealed strong UnaG fluorescence in the spheroid core, confirming oxygen depletion and formation of a hypoxic zone (Fig. 6D). This magnetically guided spheroid system therefore provides a physiologically relevant platform for modeling tumor hypoxia and was subsequently used to examine stromal assembly and functional interactions with reprogrammed fibroblasts.

### Recapitulating tumor–stromal crosstalk in hypoxic 3D co-cultures

To investigate stromal influences on tumor organization under hypoxic conditions, we established 3D co-cultures of PC-3 cells with either parental pCAFs or VDR-rpNFs. In co-cultures with VDR-rpNFs, PC-3 cells exhibited minimal association with stromal cells and adopted a diffuse, disorganized morphology (Fig. 7A–B, right panels). In contrast, pCAFs or eGFP/pCAFs promoted robust tumor–stromal contacts, with PC-3 cells forming cohesive ring-like structures around fibroblasts (Fig. 7B, left panels).

**Fig. 7.**
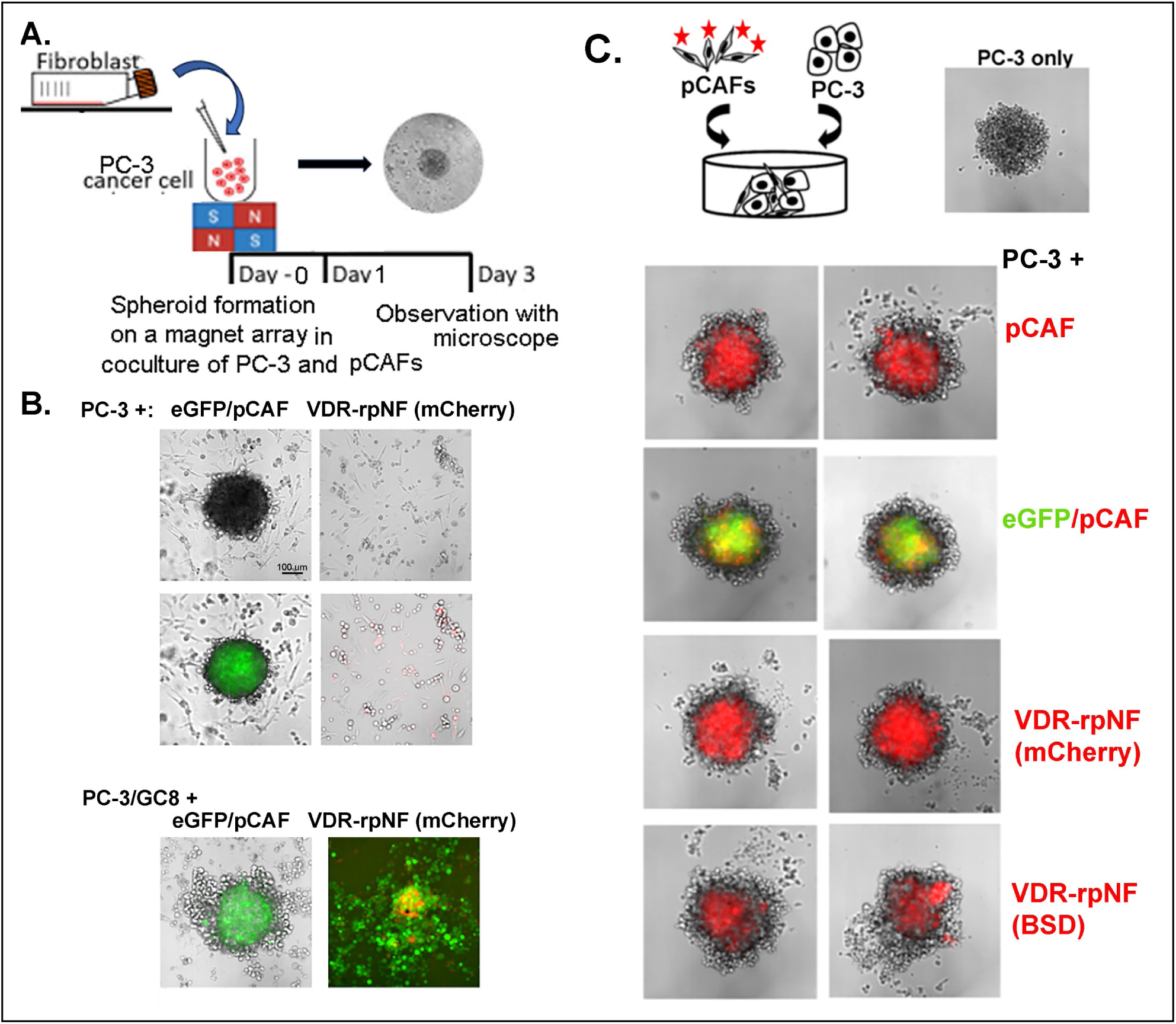
**A. B.** Coculture of 3D PC-3 or PC-3/GCamp8 with pCAFs or VDR-rpNFs (VDR-mCherry/pCAF). **C.** Reorganization of PC-3 cancer cells (unlabeled) and pCAFs or VDR-rpNFs labeled with Cellbrite Red was observed over 3 days to form macroscopic mini-tumors. One representative imaging result was shown here from at least three biological repeats.

To visualize these interactions, pCAFs or rpNFs were labeled with Cellbrite Red and incorporated into PC-3 spheroids at a 1:1 ratio (Fig. 7C). Consistent with earlier observations, VDR-rpNFs failed to support organized tumor architecture, resulting in disrupted PC-3 arrangements (bottom panels). Conversely, parental pCAFs facilitated the formation of continuous circular tumor structures (top panels), highlighting their capacity to sustain tumor–stromal crosstalk within a hypoxic 3D microenvironment.

### Interactions with other reprogrammed fibroblasts

To determine whether TFs beyond VDR similarly alter CAF behavior, we examined PPARγ-rpNFs and p53-rpNFs in 3D co-culture. Both populations showed reduced expression of canonical CAF markers (Fig. 3), indicating successful partial reprogramming. For visualization, pCAFs or rpNFs were labeled with Cellbrite Blue and co-cultured with Red-labeled PC-3 cells at a 1:1 ratio (Fig. 8).

**Fig. 8.**
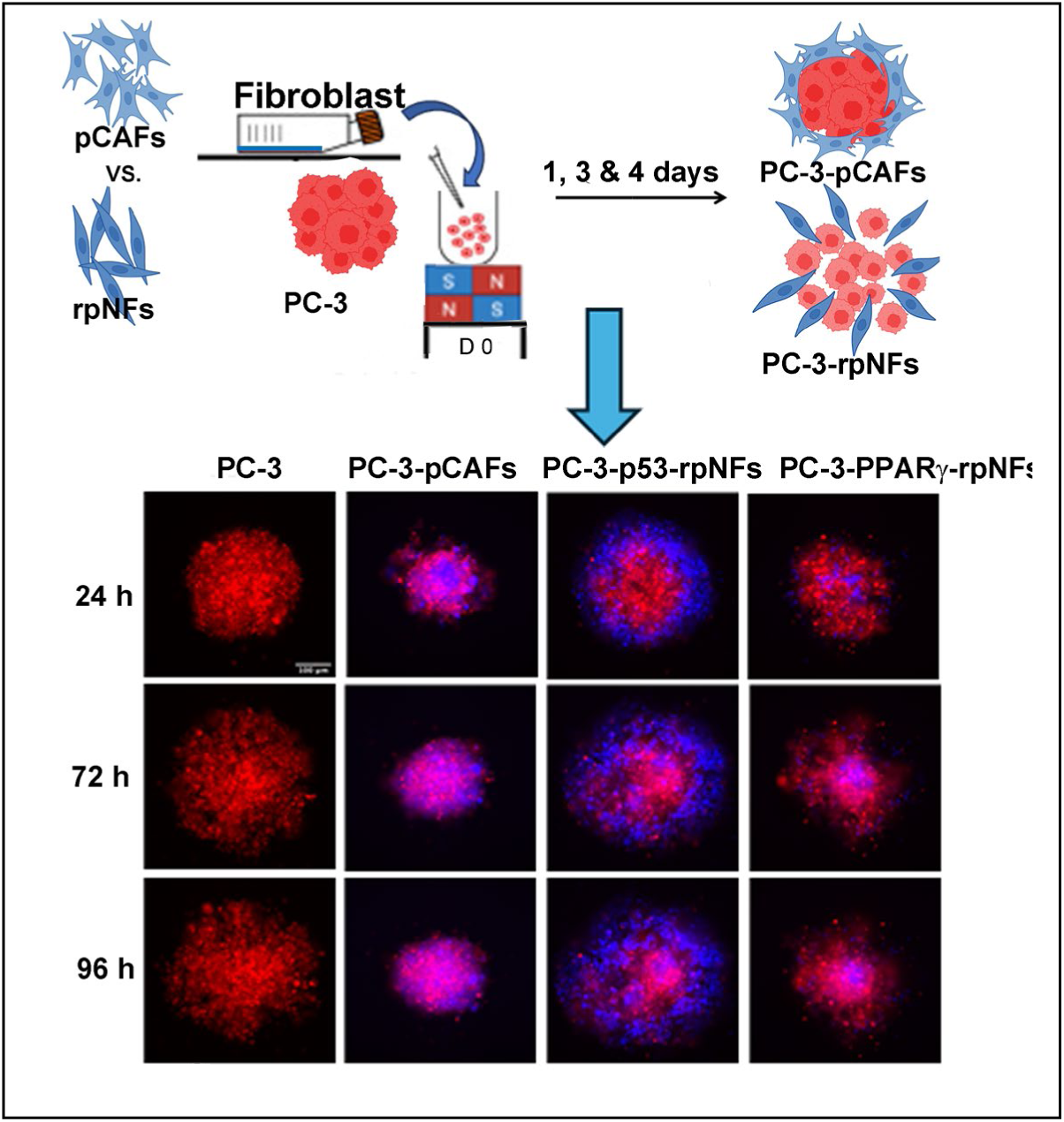
PC-3 spheroids co-cultured with parental pCAFs formed compact, cohesive structures, whereas co-cultures with PPARγ-rpNFs or p53-rpNFs produced disrupted, looser spheroids, diminishing the barrier properties of pCAFs, potentially facilitating immune cell infiltration into the tumor microenvironment.

PC-3 cells alone formed large spheroids (Fig. 8, 1st panel), whereas co-culture with parental pCAFs restricted spheroid size and promoted compact, ring-like structures (2nd panel). In contrast, co-cultures with PPARγ-rpNFs or p53-rpNFs produced disrupted, loosely organized PC-3 structures (3rd and 4th panels). These findings suggest that rpNFs generated by PPARγ or p53 transduction diminish the barrier-forming properties of pCAFs, resulting in less cohesive tumor spheroids—an architectural shift that may facilitate immune cell infiltration.

### B7-H3 expression in PC-3 and stromal cells and cytotoxicity of anti-B7-H3 CAR T cells

B7-H3 (CD276) is highly expressed on PC-3 prostate cancer cells and prostate CAFs, consistent with its roles in tumor progression and immune evasion. Immunocytochemistry confirmed strong surface B7-H3 expression on both PC-3 cells and pCAFs (Fig. 9A), supporting the rationale for B7-H3–targeted therapies capable of engaging both malignant and stromal compartments.

**Fig. 9.**
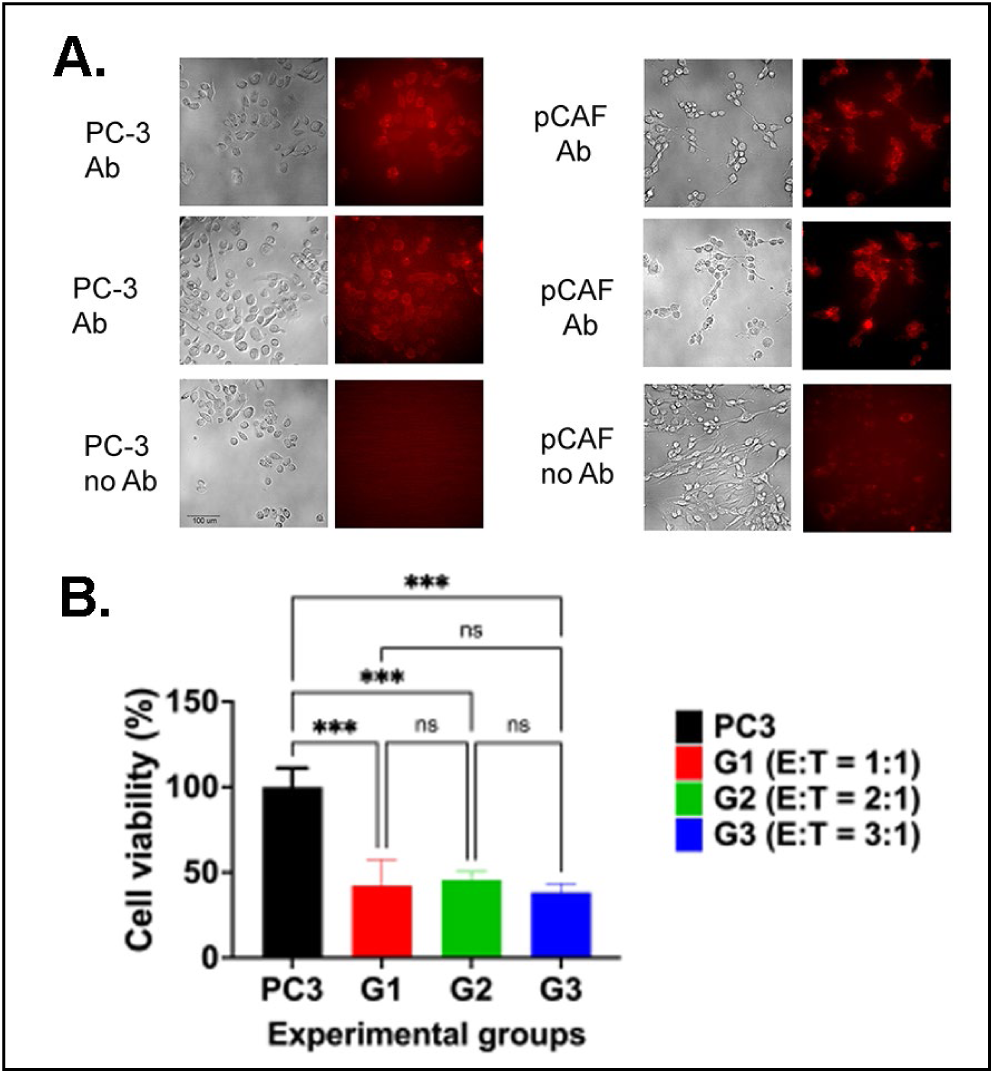
**A.** Immunocytochemistry (ICC) analysis demonstrates strong B7-H3 (CD276) expression on both PC-3 cancer cells and pCAFs, visualized by red fluorescence. **B.** In 2D co-culture assays, anti-B7-H3 CAR T cells exhibit antigen-specific cytotoxicity against PC-3 cells, with killing efficiency increasing across graded effector-to-target (E:T) ratios.

To assess anti-tumor activity, we used CD8α/CD28-B7-H3 CAR T cells engineered to express 41BBL, selected for enhanced effector function^48^. In 2D co-culture assays, B7-H3 CAR T cells exhibited robust antigen-specific cytotoxicity across all effector-to-target (E:T) ratios, with maximal killing at 3:1 (61.7 ± 15.1%; Fig. 9B). This ratio was used for subsequent 3D co-culture experiments.

### Immune Cell Infiltration in 3D Cocultures

Because PC-3 cells lack surface CD19 expression ^49^, CD19 CAR T cells—which do not kill PC-3—were used to evaluate T-cell–tumor interactions independent of cytotoxicity. mCherry-labeled CD19 CAR T cells were introduced into 3D cocultures at a 3:1 effector-to-target ratio one day after printing. In PC-3–only spheroids, T cells readily infiltrated tumor structures, as indicated by robust intratumoral red fluorescence (Fig. 10A, B). In contrast, the presence of pCAFs markedly restricted T-cell access, with fluorescence confined to the spheroid periphery, consistent with a CAF-mediated physical barrier in 3D tumor environments. Notably, VDR-rpNFs supported substantially greater T-cell penetration and tumor engagement (Fig. 10C, D, bottom). These findings demonstrate that disrupting CAF-derived structural barriers—or replacing CAFs with rpNFs—enhances immune cell access to tumor cells.

**Fig. 10.**
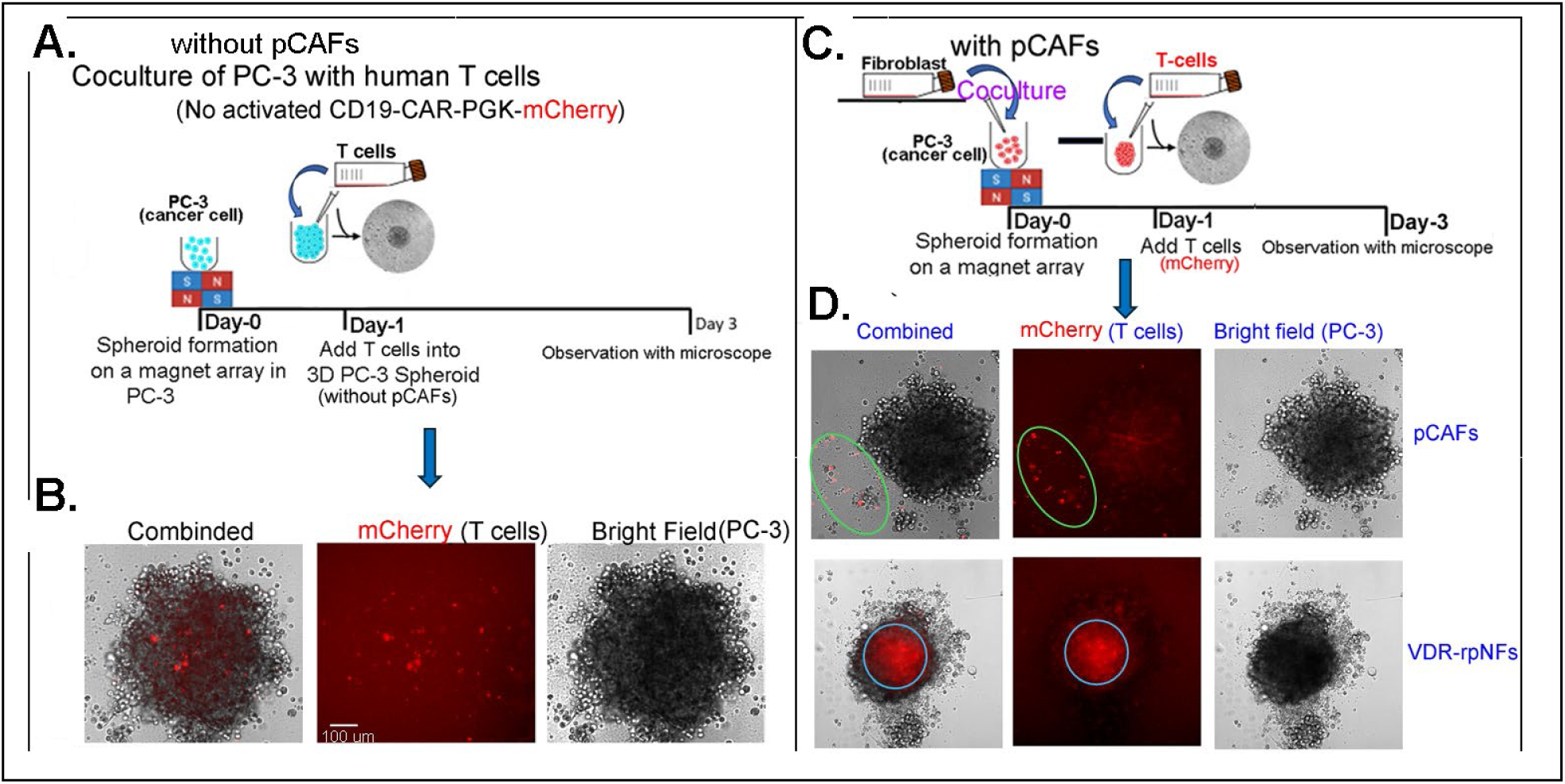
CAFs restrict T-cell interaction and infiltration in 3D prostate cancer cocultures. **(A, B)** Confocal imaging of CD19 CAR T cells (mCherry-labeled, red) added at a 3:1 effector-to-target ratio one day after printing. In the absence of pCAFs, T cells infiltrated and interacted with PC-3 cells, as indicated by red fluorescence within tumor cells. **C, D.** Coculture of 3D PC-3 with pCAFs or rpNFs, and T (red) cells. pCAFs restricted T-cell access, with red signals remaining outside tumor cells **(D, Top)**, whereas VDR-rpNFs facilitated enhanced T-cell interaction and infiltration into PC-3 cells (**D, bottom)**.

### Stromal Modulation of Anti-B7-H3 CAR T-Cell Activity in Magnetically Guided 3D-Bioprinted PC-3 Spheroids

To determine how stromal populations influence anti-B7-H3 CAR T-cell activity (Fig. 11A), magnetically guided 3D-bioprinted cocultures were generated using PC-3 tumor spheroids combined at a 1:1 ratio with either parental pCAFs or rpNFs (Fig. 11B). Anti-B7-H3 CAR T cells were added after 1–3 days at a 3:1 effector-to-target ratio. By day 5, CAR T cells induced robust tumor killing across all conditions (Fig. 11C, D). Because PC-3 cells localized to the spheroid periphery while stromal cells formed a central core, CAR T-cell access to tumor cells was not impeded, resulting in minimal differences between pCAF- and rpNF-containing spheroids. Additionally, B7-H3 expression on pCAFs rendered them direct CAR T-cell targets, further diminishing any stromal barrier effect.

**Fig. 11.**
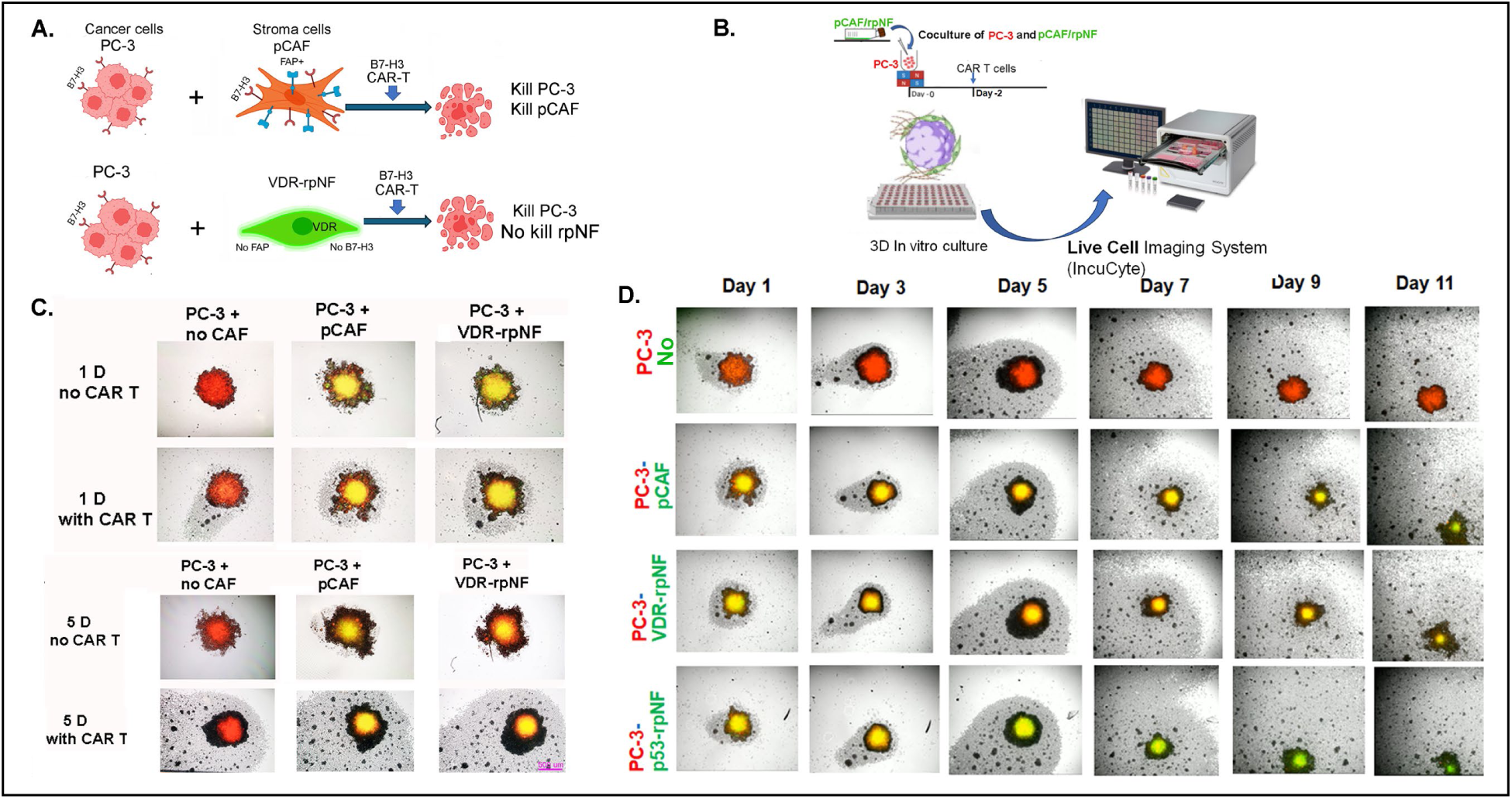
Stromal Effects on CAR T-Cell Activity in 3D-Bioprinted PC-3 Spheroids. **A.** Representative images of stromal populations influencing anti-B7-H3 CAR T-cell killing function**. B.** PC-3 spheroid with pCAF/rpNFs interaction with CAR-T cells monitored by IncuCyte imaging system. **C.** The killing effects of B7-H3 CAR T cells on PC-3 cells in day 5 compared to day 1. **D**. Imaging of live cells at different time.

Given these architectural constraints and the susceptibility of pCAFs to CAR T-cell–mediated elimination, this in vitro system was insufficient to resolve stromal contributions to CAR T-cell function, prompting evaluation in vivo.

### VDR-rpNFs Modulate Tumor Architecture and Cell-Death Programs in PC-3 Xenografts Treated With B7-H3 CAR T Cells

To investigate the role of VDR-rpNFs in shaping the prostate tumor microenvironment (TME), pilot xenograft studies were performed in male NCG mice. Animals received subcutaneous injections of 1×10⁶ PC-3-Luc cells admixed with either 1×10⁶ pCAFs or 1×10⁶ VDR-rpNFs, followed by systemic administration of 2×10⁶ B7-H3 CAR T cells on day 7 (Fig. 12A). After three weeks, tumors were harvested. Xenografts containing VDR-rpNFs were on average 24% smaller (mean 924 mm³) than those containing pCAFs (1224 mm³), consistent with known variability in PC-3 tumor growth kinetics (n=3 per group; Fig. 12B).

**Fig 12.**
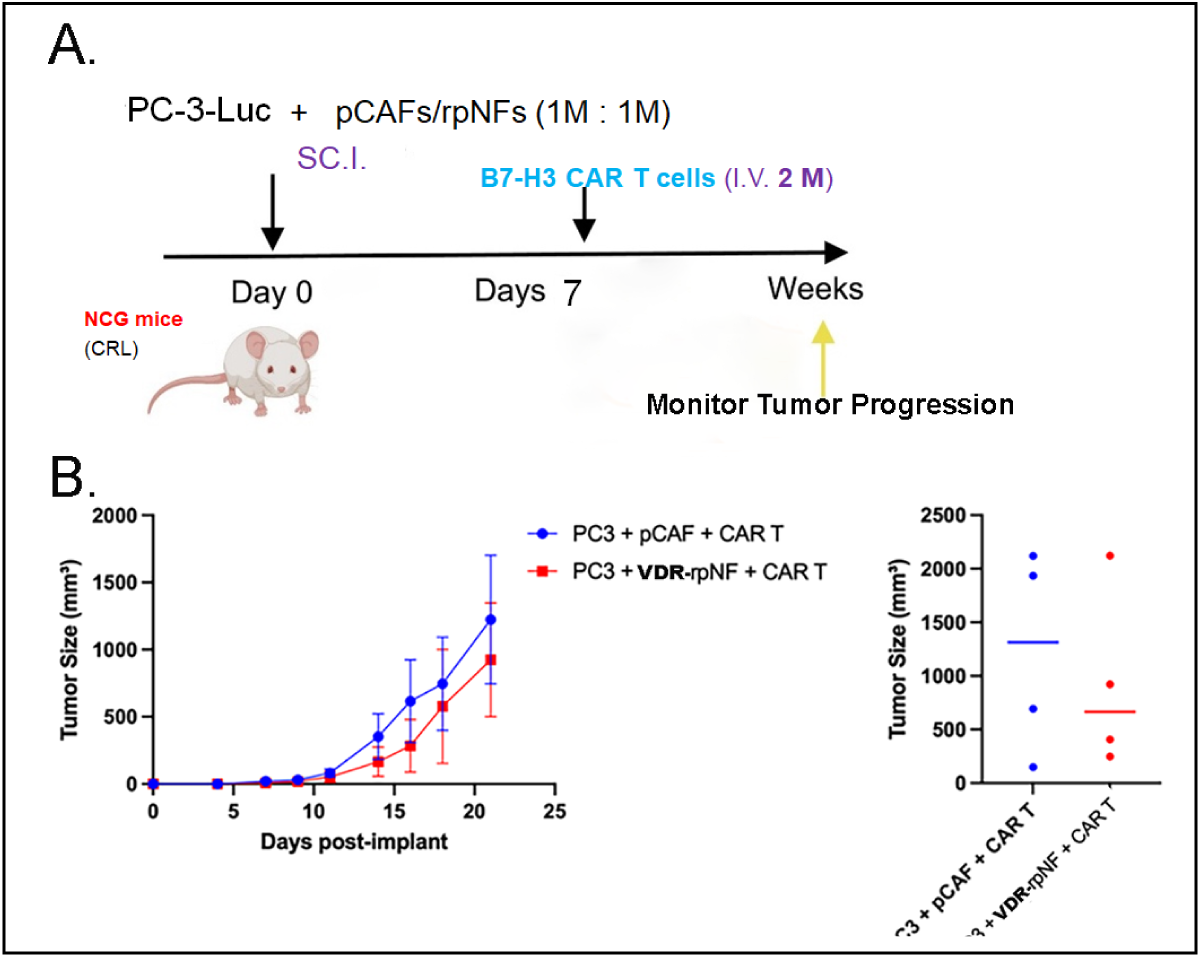
Pilot xenograft study design and tumor growth outcomes. **A.** Schematic picture of animal pilot studies. **B.** Tumor sizes of pCAF or VDR-rpNF group after injection of B7-H3 CAR T cells.

Histological analysis revealed striking differences in necrotic involvement. Tumors in the VDR-rpNF group (samples #73, #74) exhibited markedly reduced necrosis compared with pCAF-containing tumors (#68, #69; Fig. 13A). Necrosis was also less pronounced than in no-CAR-T controls (#75, #82), indicating that VDR-rpNFs confer a distinct tissue phenotype beyond both CAF-driven and untreated contexts.

**Fig. 13.**
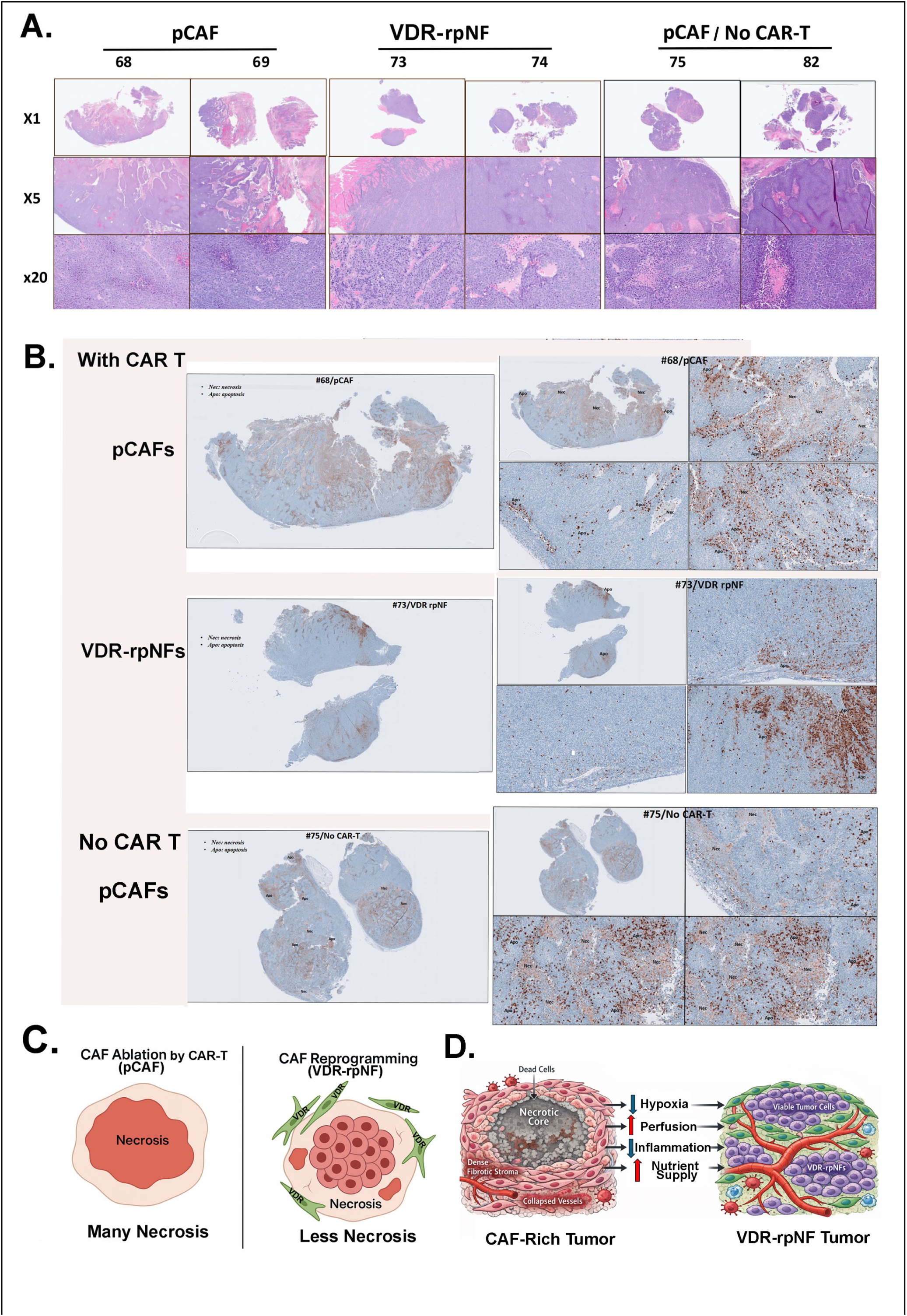
VDR-rpNFs alter tumor cell death patterns. **A.** H/E staining revealed extensive necrosis in pCAF tumors, moderate necrosis in no CAR T controls, and markedly reduced necrosis in VDR-rpNF tumors. **B.** Cleaved caspase-3 staining confirmed minimal necrosis in VDR-rpNF tumors, with apoptosis localized to the periphery, contrasting with central necrosis in pCAF tumors and moderate necrosis in no CAR T controls. **C.** Comparison of reprogramming CAFs with CAF ablation by CAR-T on Necrosis. **D.** Potential Mechanisms of necrosis reduction by VDR-rpNFs. Schematic showing that CAF-rich tumors develop dense stroma, collapsed vessels, and hypoxia driven by TGF-β and IL6, resulting in necrosis. VDR-rpNFs suppress CAF programs, restore ATP, and reduce TGF-β/IL6 signaling, improving perfusion and lowering inflammation. These changes preserve tissue viability and support CAR T-cell infiltration, thereby limiting necrosis.

Immunohistochemical staining for cleaved caspase-3 (CST D175) corroborated these findings (Fig. 13B). pCAF-treated tumors displayed extensive central necrosis, characteristic of widespread non-apoptotic cell death, whereas no-CAR-T controls showed moderate necrotic involvement. In contrast, VDR-rpNF–treated tumors exhibited minimal necrosis, with apoptosis largely confined to the tumor periphery. This spatially restricted apoptotic pattern suggests that VDR-rpNFs redirect cell-death programs away from uncontrolled necrosis toward regulated peripheral apoptosis.

Together, these results demonstrate that VDR-rpNFs substantially reduce necrosis and reshape tumor cell-death architecture in vivo. This coordinated shift may enhance immune accessibility and mitigate the pro-tumorigenic consequences of CAF-driven necrotic remodeling.

### VDR-rpNFs and CAR T Cells Independently Reduce FAP^+^ CAF Abundance

Quantitative analysis of PC-3 xenografts revealed distinct patterns of FAP^+^ CAF abundance across treatment groups (Fig. 14). No-CAR-T controls (#75) exhibited high levels of FAP^+^ CAFs, reflecting persistent fibroblast activation in the absence of targeted therapy (Fig. 14A, D). In contrast, pCAF tumors (#68), which express high levels of B7-H3, contained markedly fewer FAP^+^ CAFs (Fig. 14B, E), consistent with direct CAR T-cell–mediated elimination of B7-H3^+^ pCAFs. Strikingly, VDR-rpNF tumors (#73) showed almost no detectable FAP^+^ CAFs (Fig. 14C, F), indicating that transcription factor–mediated reprogramming effectively suppresses CAF activation. These histological observations align with PCR-based evidence of reduced FAP expression in VDR-rpNFs (Fig. 2E).

**Fig. 14.**
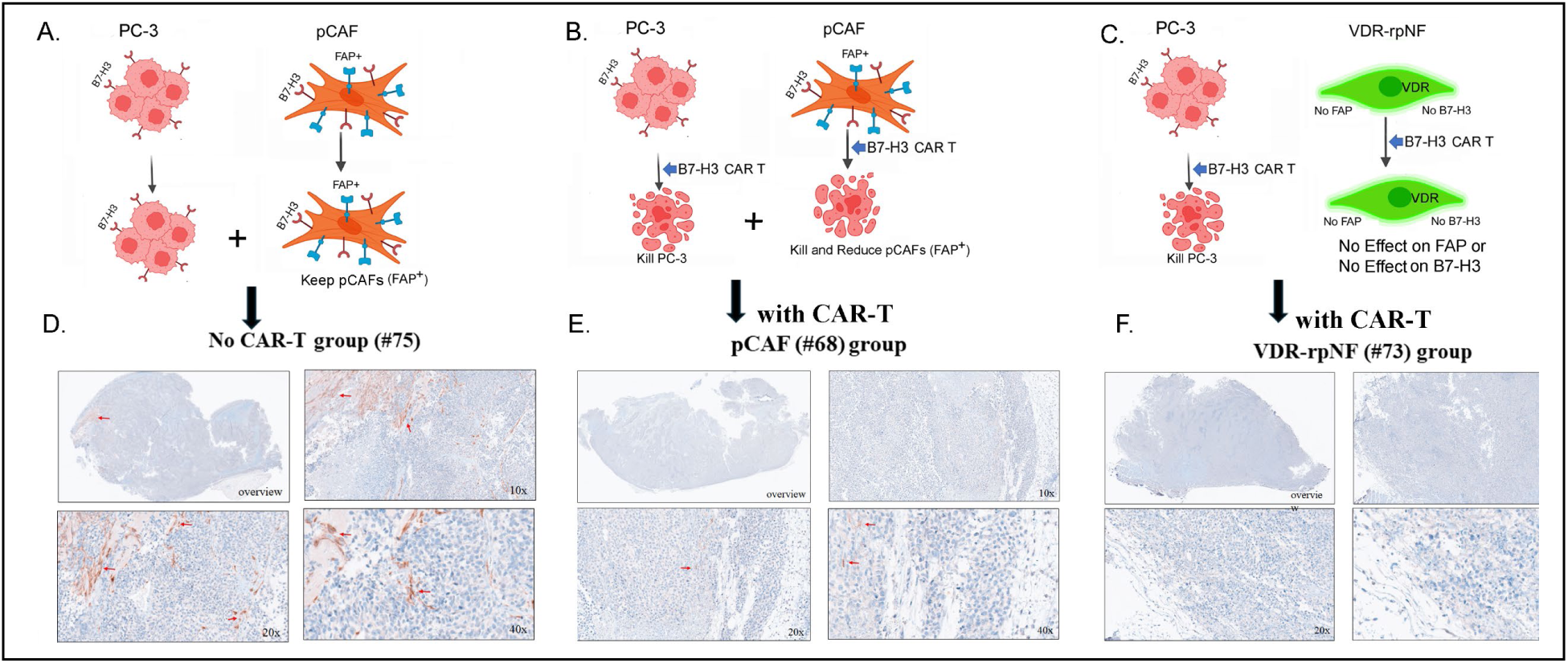
Schematic pictures of PC-3 xenograft tissues with stroma and CAR T cells. **(A-C)**, and Quantitative analysis of FAP protein expression in PC3 tumor tissues with pCAFs with no CAR-T control group (#75) **(D)**, pCAFs with B7-H3 CAR-T group (#68) **(E)** and VDR-rpNF with B7-H3 CAR-T group (#73) **(F)**.

Collectively, these findings demonstrate that both CAR T-cell cytotoxicity and VDR-driven fibroblast reprogramming diminish FAP^+^ CAF prevalence through distinct mechanisms. Together, they contribute to stromal normalization in prostate cancer xenografts and highlight the complementary therapeutic potential of combining immune-based cytotoxicity with fibroblast reprogramming.

## DISCUSSION

Our findings demonstrate that transcription factor–mediated reprogramming can partially reverse CAF identity and remodel the tumor microenvironment in ways that support immune activity. Introducing VDR, PPARγ, or p53 individually into pCAFs consistently reduced canonical CAF activation markers, indicating that pCAFs retain a degree of plasticity and can be shifted toward a more quiescent, fibroblast-like state. Although these single-factor interventions represent proof-of-concept perturbations, they parallel prior work showing that defined transcription factor combinations—such as the Yamanaka factors for iPSC induction^50,51^ or TF cocktails that convert tumor cells into cDC1-like antigen-presenting cells^52^—can drive more complete lineage reprogramming. Building on this rationale, we leveraged a transcription factor network enriched in quiescent, anti-inflammatory fibroblasts—p53, VDR, PPARγ, and RXR (53VPR)—to redirect CAFs toward a tumor-suppressive state. VDR and p53 synergistically activate tumor-suppressive gene programs, while RXR’s heterodimeric interactions with VDR and PPARγ provide a framework for coordinated transcriptional rewiring. This integrated TF architecture offers a mechanistic basis for redirecting stromal support away from tumor progression, and future studies will validate pCAF reversion using the 53VPR set through integrated multi-omics and functional assays.

Using a magnetically assisted bioprinted 3D spheroid model with a hypoxia reporter ^40–42^, we previously established a platform that recapitulates the hypoxic tumor core and CAF-driven architectural constraints. Within this system, VDR-rpNFs disrupted the dense, ring-like structures formed by parental pCAFs and enhanced PC-3 cell engagement with non-specific CAR T cells, promoting immune cell infiltration. PPARγ-rpNFs and p53-rpNFs similarly reduced CAF markers and generated loosely organized spheroids, supporting the conclusion that TF-mediated reprogramming diminishes structural barriers to immune entry. These observations reinforce the hypothesis that CAF reprogramming increases tumor permeability to immune effectors and justify further mechanistic investigation.

Pilot PC-3 xenograft studies revealed an additional role for VDR-rpNFs in modulating tumor cell death dynamics. VDR-rpNFs attenuated both necrotic and apoptotic cell death, preserving tissue integrity under CAR T–associated stress. By limiting central necrosis and reducing peripheral caspase-3 activation, VDR-rpNFs appear to dampen collateral tissue injury, which may in turn influence immune infiltration and therapeutic efficacy. These protective effects likely arise from VDR-dependent recalibration of fibroblast–immune interactions: VDR signaling modulates cytokine production, extracellular matrix pathways, and cellular stress responses^53–55^, collectively generating a microenvironment less prone to inflammatory amplification during immunotherapy. Reduced release of necrosis-associated DAMPs may further limit secondary immune activation, while decreased caspase-3 activity suggests altered paracrine signaling or enhanced survival cues within adjacent tumor cells.

Necrosis is a hallmark of solid tumors in which proliferative demand exceeds vascular supply^56^, and pCAF-supported xenografts displayed prominent central necrosis consistent with CAF-driven hypoxia and nutrient deprivation^57^. In contrast, VDR-rpNF tumors were smaller and exhibited markedly reduced necrotic cores, indicating that stromal reprogramming not only restrains tumor expansion but also mitigates the microenvironmental stresses that precipitate necrotic collapse. Reduced necrosis has several mechanistic implications: improved perfusion and oxygenation align with stromal normalization^58^ (Fig. 13 C,D); lower interstitial pressure may enhance nutrient, oxygen, and therapeutic diffusion; diminished necrotic debris may alleviate immunosuppressive inflammation driven by DAMP-mediated recruitment of suppressive myeloid cells^59^; and preserved architecture may improve responsiveness to immune-based therapies, which are often hindered by necrotic barriers and disorganized stroma^60^.

Together, these findings indicate that VDR-mediated CAF reprogramming reshapes fibroblast–immune interactions in ways that stabilize the tumor microenvironment during CAR T-cell therapy. By dampening inflammatory signaling, reducing DAMP release, and modulating apoptotic pathways, VDR-rpNFs limit necrotic and apoptotic tissue damage while preserving stromal structure. This controlled remodeling contrasts with the extensive tissue disruption observed after CAF ablation and suggests that engineering fibroblast behavior—rather than eliminating the stromal compartment—may broaden the therapeutic window of adoptive immunotherapy. As fibroblasts represent one of the most abundant and influential stromal populations in solid tumors, the ability to reprogram their function offers a conceptual shift in how the microenvironment can be leveraged to enhance treatment efficacy. Future work integrating rational TF combinations, multi-omics profiling, and more reproducible xenograft systems will be essential for defining the full potential of fibroblast engineering as a complementary modality in solid tumor immunotherapy.

## ACKNOWLEDGEMENT

This work was supported in part by grants from the National Institutes of Health (NIH) EB029122, GM140929, and CA262815 to Y. Wang, and by a private Salvaterra family foundation to J. Pinski. This work was also supported by the USC Preclinical Models Shared Resource (funded by Norris Comprehensive Cancer Center CCSG grant, NCI grant # P30CA014089). The graphical abstract was modified from a graphic created with BioRender.com.

## AUTHOR CONTRIBUTIONS

Conception and design: N.S.L. Acquisition of data: N.S.L., P.D., Y.H., B.R., X.Y., T.G, P.H. Development of methodology: N.S.L, Y.H., P.D. Acquisition of data in 3D printing: P.D., S.M. Development and culture for B7-H8 CAR T cells: C.W.Y. and C.DR. Acquisition of data in *in vivo* analysis: B.Y.T, S.X. Analysis and interpretation of data: N.S.L, P.D., Y.H., S.X. Writing, review, and/or revision of manuscript: all authors. Study supervision: N.S.L., J.P. and I.K.P.

## METHODS

### Cell culture

All cells are cultured in 5 % CO2 at 37°C. Prostate CAFs (pCAFs) are acquired from American Type Culture and Collection (ATCC, hTERT PF179T) and maintained in the complete DMEM medium supplemented with 10% FBS, 7.5% sodium bicarbonate and 1 µg/mL Puromycin. HEK293T and PC-3 cells are maintained in the complete DMEM medium supplemented with 10% FBS and 1% Penicillin-Streptomycin (P/S). HDFs are maintained in the complete DMEM supplemented with 20% FBS and 1% P/S.

### Construction of pCAFs containing TFs

Prostate CAFs (pCAFs; hTERT PF179T) are purchased from ATCC and used, and Normal human dermal fibroblasts (HDF) are used for NFs as a control. In order to get high induction of pCAFs, lentiviral vectors with high transduction are used. 1 day prior to transfection, pCAF cells are seeded at ∼40% confluency in 6- or 12-well plates. Cells are transfected the next day at ∼80% confluency. Plasmids containing the lentivector of interest (VDR, PPARg or p53) (Addgene plasmid # 145005, TFORF3529 (VDR); TFORF 3139 (PPARγ); pLenti6/V5-p53_wt p53) are purchased from Addgene. After removing the Puromycin gene from the vector by using KpnI/HpaI cut and insert PCR products of Blasticidinr or mCherry genes containing KpnI/HpaI sites. Each plasmid, psPAX2, and pMD2.G are transfected using Lipofectamine 3000 and P3000 Enhancer. Media is changed 5 h after transfection. Virus supernatant is harvested 48 h post-transfection, centrifuged and filtered with a 0.45 mm PVDF filter. The filtered virus is concentrated with Lenti-X concentrator to get higher titer, aliquoted, and stored at -80°C.

We used two different methods for selecting the transduced pCAFs: Antibiotic and Fluorescence. For selecting pCAFs transduced by the lentivirus containing VDR-Blasticidin^r^ genes, 50% density of pCAFs are seeded in 12-well plates with an appropriate volume of lentivirus. After 24h, media is refreshed with the appropriate antibiotic (2 or 3 μg/mL Blasticidin). For 5 days, media with the appropriate antibiotic is refreshed every day, and cells are passaged after 3 days of selection (‘pCAF/VDR-BSD’). The parental pCAFs (no transduction) are dead at 1 day after addition of 2 or 3μg/mL Blasticidin. Another selection method is to use fluorescence activated cell sorting (FACS). pCAFs transduced by lentivirus containing eGFP (control, ‘pCAF/eGFP’) or VDR-mCherry (‘pCAF/VDR-mCherry’) are selected by FACS).

### Q-RT-PCR and FACS assays

#### Q-RT-PCR

To assess the expression of specific genes in transduced pCAF cells, RNA purification and cDNA synthesis is performed. Total RNA is isolated from different CAFs using QuickRNA Miniprep (Zymo Research). First strand cDNA is synthesized using iScript cDNA synthesis kit (Bio-Rad) according to the manufacturer’s instructions. Samples are run in 10 ul reactions using an CFX384 System (Bio-Rad). SYBR Green oligonucleotides will be used for detection and quantification of genes. The primers for detectable genes are synthesized by Integrated DNA Technologies (IDT). Gene expression levels will be calculated after normalization to the standard housekeeping gene GAPDH or Actin using Confidential the ΔΔ CT method and expressed as relative mRNA levels compared with control. All primer sequences are listed in the Supplementary section.

#### FACS assay

Single cell suspensions are stained with the indicated antibodies. Briefly, cells are pelleted, resuspended in the staining solution with fluorescent antibodies, and incubated for 1 hr at 4°C in the dark. Next, the cells are washed and resuspended in PBS at an approximate density of 5 × 10^6^ cells/mL. CD44 as a cancer-associated fibroblast cell surface marker (29) and CD39 as a marker for normal fibroblasts from healthy tissues are used. Their antibodies conjugated to fluorescent dyes are used to detect the proteins expressed in pCAFs and rpNFs. FACS analysis is performed using an LSR II flow cytometer (BD Biosciences) and analysed using FlowJo. For FACS analysis, CAFs are stained with phycoerythrin (PE)-conjugated anti-CD39, or anti-CD44 and analyzed by flow cytometry analyzer (Excitation: 488-561 nm; Emission: 578 nm).

### Western blot analysis of VDR expression in VDR-rpNFs or pCAFs

VDR protein expression was evaluated in control pCAFs and VDR-rpNFs by Western blotting. Cells were washed with ice-cold PBS and lysed in RIPA buffer supplemented with protease and phosphatase inhibitors on ice for 30 min, followed by clarification at 14,000 × g for 15 min at 4°C. Protein concentration was determined by BCA assay, and equal amounts of protein (typically 20–30 µg per lane) were mixed with 4× Laemmli sample buffer containing β-mercaptoethanol, boiled at 95°C for 5 min, and resolved on 8–10% SDS–polyacrylamide gels. Proteins were transferred to PVDF membranes, which were then blocked in 5% non-fat dry milk in TBST for 1 h at room temperature and incubated with primary antibody against human VDR (R&D Systems, catalog number PP-H4537-00, 1/1,000 dilution) overnight at 4°C. After washing in TBST, membranes were incubated with HRP-conjugated secondary Goat anti-Mouse antibodies (Abcam, 1/2,000 dil.) for 1 h at room temperature and developed using enhanced chemiluminescence. Band intensities were quantified using ImageJ and normalized to β-actin as a loading control. Relative VDR expression in VDR-rpNFs was expressed as fold change over control pCAFs.

### Spheroid printing

Cell spheroids are printed using a high-throughput, magnetic field-guided, labelfree method. Briefly, PC-3 cells are seeded in an uncoated, flat, clear bottom, square 96-well plate (ibidi, USA) with 5000 cells/well in a culture medium containing a paramagnetic salt, Gadavist. Based on our previous works the concentration of Gadavist in the culture medium is maintained at 25 mM. The well plate is placed on an inhouse designed magnetic array containing cuboid permanent magnets (N52-grade neodymium has dimensions 4.5 mm × 4.5 mm × 4.5 mm). The detailed protocol is in our previous papers. This magnetic force ultimately is driving the PC-3 cells to form 3D structures at the lowest magnetic field zone within 3 hours.

### ATP release assay

ATP release is assessed after printing of the spheroids at regular intervals (1 day, 2 days, and 4 days) using the Promega CellTiter-Glo® 3D cell viability assay kit according to the manufacturer’s protocol. Briefly, following the spheroid printing and incubation, the old medium is aspirated, and the assay reagent is added to the fresh culture medium at a 1:1 (v/v) ratio in each well of the 96-well plates (ibidi 89621) and mixed vigorously for 5 min to induce cell lysis. After mixing, the plate is kept at room temperature for 25 min, and a dark condition is ensured. Finally, the supernatant is collected into a white opaque tissue culture-treated 96-well plate (353296, Becton Dickinson Labware, USA) to minimize signal loss. The luminescence is measured (integration time 0.01 s) using a plate reader (Synergy H4). The data is processed and analyzed using the GraphPad Prism (version 10) software.

### Live-cell Immunocytochemistry (ICC) for B7-H3 protein expression

Live-cell staining was performed to assess surface expression of B7-H3 (CD276). Cells were washed gently with cold PBS and incubated in PBS containing 1–2% BSA for 10–15 min at 4 °C to reduce nonspecific binding. Primary human anti–B7-H3 antibody (AF1027, R&D Systems) was diluted in cold staining buffer (0.5 µg/µL; 2 µL per 100 µL reaction volume) and applied to cells for 20–30 min at 4 °C. After incubation, cells were washed 2–3 times with cold PBS using minimal pipetting force to preserve cell attachment. When required, fluorophore-conjugated secondary antibodies (α-mouse-Alexa Fluor 594, red) were added for 20 min at 4 °C, followed by gentle washing. Cells were imaged immediately in live-cell imaging medium or cold PBS by a fluorescence microscopy as fluorescence signal stability decreases over time.

### Primary T cell isolation and transduction of B7-H3 CAR T construct

Peripheral blood mononuclear cells (PBMCs) were obtained from buffy coats (Excellos) and isolated using lymphocyte separation medium (Corning, 25-072-CV) according to the manufacturer’s instructions. Primary human T cells were enriched from PBMCs using the Pan T Cell Isolation Kit (Miltenyi, 130-096-535) and activated the same day with Dynabeads Human T-Expander CD3/CD28 (Gibco, 11141D). On day 3 post-activation, Dynabeads were magnetically removed, and cells were transduced with concentrated lentiviral particles at a multiplicity of infection (MOI) of 10 in 24-well plates coated with RetroNectin® (30 µg/mL; Takara, T100B). Plates were centrifuged at 1,800 × g for 1 h at 32 °C and then returned to standard culture conditions. For enrichment of CAR-expressing cells, transduced T cells were stained with Alexa Fluor® 647 AffiniPure F(ab′)_2_ Fragment Goat Anti-Human IgG, F(ab′)_2_ fragment specific (Jackson ImmunoResearch, 109-606-097) and sorted on a Sony SH800 cell sorter to isolate a pure CAR^+^ population for downstream applications.

### Cytotoxicity of B7-H3 CART cells on PC-3 tumor cells in 2D and 3D experiments

To determine the killing efficacy of the B7-H3 CART cells on PC3 tumor cells, firstly, 2D experiments are carried out. For this purpose, PC3 cells are seeded (5000 cells/well) in a tissue culture-treated 96-well plate. After 48 hrs., the old media is replaced with fresh culture media, and the B7-H3 CART cells are added at three different effectors to target ratios (1:1, 2:1, and 3:1) into the treatment groups. The cells are co-cultured for 5 days, and an endpoint cell viability assay is performed using the CCK8 kit following the manufacturer’s protocol. After co-culturing, the old medium is aspirated, and 100 µL of fresh medium is added. 10 µL of CCK-8 solution is added to each well and incubated for 1 hr. Finally, absorbance is measured using a plate reader (Synergy H4) at 450 nm, and the data is post-processed using the GraphPad Prism (version 10) software.

### Animal study

All animal experiments were conducted in accordance with Institutional Animal Care and Use Committee guidelines and an approved protocol. Male 6-week old NCG mice (Charles River Laboratory) were injected subcutaneously in the right flank with 1x10^6^ PC-3-Luc mixed with 1x10^6^ pCAFs or 1x10^6^ rpNFs (n=5 per group). Tumor volume was calculated using length × width × width × 0.5. 2x10^6^ CAR T cells were injected intravenously after 7 days. Mice were euthanized at the study endpoint on day 21. Tumors were harvested, and half was preserved in 10% formalin, while the other half was flash frozen. The lung, liver, spleen were also harvested and preserved in 10% formalin.

### Statistical Analyses

In general results are expressed as mean ± s.e.m. Statistical analysis of multiple groups used one-way ANOVA, and Dunnet’s correction for multiple comparisons (GraphPad Prism, V8). Two group comparisons were tested using the Student’s t-test (one and two-tailed) in Excel (v2016).

## REFERENCES

1. Kuttiappan, A., Chenchula, S., Vardhan, K.V., et al. 2025. CAR T-cell therapy in hematologic and solid malignancies: mechanisms, clinical applications, and future directions. Medical Oncology 42:376.

2. Guo, J., Zhou, C., Zhao, H., Li, H. 2025. Challenges and breakthroughs: current landscape and future prospects of CAR-T cell therapy clinical trials for solid tumors. Frontiers in Oncology 15:1652329.

3. Mahat, U., Das, A., Kumar, V., Prasad, V. 2025. Advancing CAR T-Cell Therapy: Evidence-Based Trial Design for Chimeric Antigen Receptor T-Cell Therapy in Oncology. JAMA.

4. Huang, Z., Chen, J., Zhu, T. et al. 2025. Cancer-associated fibroblasts in the tumor microenvironment: heterogeneity, crosstalk mechanisms, and therapeutic implications. Mol Cancer

5. Ruan, G., Wang, X., Ou, H. & Guo, D. 2025. Cancer-associated fibroblasts: dual roles from senescence sentinels to death regulators and new dimensions in therapy. Frontiers in Immunology (Sec. Cancer Immunity and Immunotherapy).

6. Hilmi M, Nicolle R, Bousquet C, Neuzillet C. 2020. Cancer-Associated Fibroblasts: Accomplices in the Tumor Immune Evasion. Cancers (Basel) 12(10):2969.

7. Pawar, J.S., Salam, M.A., Dipto, M.S.U., Al-Amin, M.Y., Salam, M.T., Sengupta, S., Kumari, S., Gujjari, L., Yadagiri, G. 2025. Cancer-Associated Fibroblasts: Immunosuppressive Crosstalk with Tumor-Infiltrating Immune Cells and Implications for Therapeutic Resistance. Cancers 17, 2484.

8. Lan, X., Li, W., Zhao, K., Wang, J., Li, S., Zhao, H. 2025. Revisiting the role of cancer-associated fibroblasts in tumor microenvironment. Front. Immunol., Sec. Cancer Immunity and Immunotherapy 16.

9. Ping, Q., Yan, R., Cheng, X., et al. 2021, Cancer-associated fibroblasts: overview, progress, challenges, and directions. Cancer Gene Therapy 28:984–999.

10. Lan, X., Li, W., Zhao, K., et al. 2025. Revisiting the role of cancer-associated fibroblasts in tumor microenvironment. Frontiers in Immunology 16.

11. Kisseleva, T., Cong, M., Paik, Y., Scholten, D., Jiang, C., Benner, C., Iwaisako, K., Moore-Morris, T., Scott, B., Tsukamoto, H., Evans, S.M., Dillmann, W., Glass, C.K., Brenner, D.A. 2012. Myofibroblasts revert to an inactive phenotype during regression of liver fibrosis. Proc Natl Acad Sci U S A 109(24):9448–53.

12. Mayorca-Guiliani, A.E., Leeming, D.J., Henriksen, K., et al. 2025. ECM formation and degradation during fibrosis, repair, and regeneration. Nature Reviews Metabolic health and disease 3:25.

13. Weiskirchen, R. 2025. Exploring Molecular Mechanisms of Liver Fibrosis. International Journal of Molecular Sciences 26(1):326.

14. Talbott, H.E., et al. 2022. Wound healing, fibroblast heterogeneity, and fibrosis Cell Stem Cell 29 (8): 1161 – 1180.

15. Younesi, F.S., Miller, A.E., Barker, T.H., Rossi, F.M.V., Hinz, B. 2024. Fibroblast and myofibroblast activation in normal tissue repair and fibrosis. Nature Reviews Molecular Cell Biology 29 (8): 1161–1180.

16. Roman, J. 2023. Fibroblasts—Warriors at the Intersection of Wound Healing and Disrepair. Biomolecules 13(6):945.

17. Sherman, M.H., et al., 2014. Vitamin D Receptor-Mediated Stromal Reprogramming Suppresses Pancreatitis and Enhances Pancreatic Cancer Therapy. Cell 159, 80–93.

18. Evans, R.M. 2014. Vitamin D receptor stromal reprogramming suppresses pancreatitis and enhances pancreatic cancer therapy. Cancer Res 74 (19): SY04–04.

19. Grosu, I., Constantinescu, A., Balta, M.D., et al. 2024.Vitamin D and Pancreatic Ductal Adenocarcinoma—A Review of a Complicated Relationship. Nutrients 16(23):4085.

20. Rowley, D.R. 2014. Reprogramming the Tumor Stroma: A New Paradigm. Cancer Cell 26 (4): 451 – 452.

21. Wang, Z., Dong, S., Zhou, W. 2024. Pancreatic stellate cells: Key players in pancreatic health and diseases. Mol Med Rep 30: 109.

22. Litewka, J.J., Jakubowska, M.A., Targosz-Korecka, M., et al. 2025. Biophysical and biochemical signatures of pancreatic stellate cell activation: insights into mechano-metabolic signalling. Cell Communication and Signaling 23:363.

23. Villapol, S. 2018. Roles of Peroxisome Proliferator-Activated Receptor Gamma on Brain and Peripheral Inflammation. Cell Mol Neurobiol 38(1):121–132.

24. Hazra, S., Xiong, S., Wang, J., Rippe, R.A., Chatterjee, V.K.K. and Tsukamoto. H. 2004. Peroxisome Proliferator-activated Receptor γ Induces a Phenotypic Switch from Activated to Quiescent Hepatic Stellate Cells. J. Biol Chem 279:11392–11401.

25. Masamune, A., Kikuta, K., Satoh, M., Sakai, Y., Satoh, A. and Shimosegawa, T. 2002. Ligands of Peroxisome Proliferator-activated Receptor-Block Activation of Pancreatic Stellate Cells. J. Biol Chem 277:141–147.

26. Aubrey, B.J., Kelly, G.L., Janic, A., Herold, M.J. and Strasser, A. 2018. How does p53 induce apoptosis and how does this relate to p53-mediated tumour suppression? Cell Death Differ 25(1):104–113.

27. Thomas, A.F., Kelly, G.L. and Strasser, A. 2022. Of the many cellular responses activated by TP53, which ones are critical for tumour suppression? Cell Death Differ 29(5):961–971.

28. Öhlund, D., Elyada, E., Tuveson, D. 2014. Fibroblast heterogeneity in the cancer wound. Journal of Experimental Medicine 211(8):1503–1523.

29. Lan, X., Li, W., Zhao, K., et al. 2025. Revisiting the role of cancer-associated fibroblasts in tumor microenvironment. Frontiers in Immunology 16:1582532.

30. Saxena, M., van der Burg, S.H., Melief, C.J.M., Bhardwaj, N. 2021. Therapeutic cancer vaccines: from design to clinical application. Nature Reviews Cancern 21:360–378.

31. Hinz, B., Lagares, D. 2020.Fibroblasts in fibrosis: novel roles and therapeutic targets. Nature Reviews Drug Discovery 19(9):699–717.

32. Agorku, D.J., Langhammer, A., Heider, U., Wild, S., Bosio, A., Hardt, O. 2019. CD49b, CD87, and CD95 Are Markers for Activated Cancer-Associated Fibroblasts Whereas CD39 Marks Quiescent Normal Fibroblasts in Murine Tumor Models. Frontiers in Oncology 9:716.

33. Faubert, B., Solmonson, A., DeBerardinis, R.J. 2020. Metabolic reprogramming and cancer progression. Science 368(6487):eaaw5473.

34. Vander Heiden, M.G., Cantley, L.C., Thompson, C.B. 2009. Understanding the Warburg effect: the metabolic requirements of cell proliferation. Science 324(5930):1029–1033.

35. Yang, J., Xu, T. & Xu, R. 2026. Vitamin D receptor suppresses pulmonary fibroblast activation by downregulating the TGF-β1/Smad signaling pathway. Histochem Cell Biol 164, 12.

36. Young, et al., 2025. IL-6 underlies microenvironment immunosuppression and resistance to therapy in glioblastoma. bioRxiv, 2025

37. Wu, S., Cao, Z., Lu, R., Zhang, Z., Sethi, G., You, Y. 2025. Interleukin-6 (IL-6)-associated tumor microenvironment remodelling and cancer immunotherapy Cytokine & Growth Factor Reviews 85, 93–102.

38. Matsuda, Satoru et al. 2023. TGF-β in the microenvironment induces a physiologically occurring immune-suppressive senescent state. Cell Reports, 42 (3): 112129

39. Perez-Penco, M., Byrdal, M., et al., 2025. The antitumor activity of TGFβ-specific T cells is dependent on IL-6 signaling. Cellular & Molecular Immunology 22:111–126.

40. Mishriki, S., Aithal, S., Gupta, T., Sahu, R.P., Geng, F. and Puri, I.K.. 2020. Fibroblasts Accelerate Formation and Improve Reproducibility of 3D Cellular Structures Printed with Magnetic Assistance. Research (Wash DC) 2020:3970530.

41. Gupta, T., Sahu, R.P., Dabaghi, M., Zhong, L.S., Shargall, T., Hirota, J.A., Carl D. Richards, C.D. and Puri, I.K. 2023. Biophysical and Biochemical Regulation of Cell Dynamics in Magnetically Assembled Cellular Structures. ACS Omega 8, 19976.

42. Datta, P., Lee, N.S., Moolayadukkam, S., Sahu, R.P., Yu, X., Guo, T., Zhou, Q., Wang, Y and K. Puri, I.K. 2024. In Vitro Sonodynamic Therapy Using a High Throughput 3D Glioblastoma Spheroid Model with 5-ALA and TMZ Sonosensitizers. Adv. Healthcare Mater. 2402877.

43. Erapaneedi, R., Belousov, V.V., Schäfers, M., Kiefer, F. 2016. A novel family of fluorescent hypoxia sensors reveal strong heterogeneity in tumor hypoxia at the cellular level. EMBO Journal 35(1):102–113.

44. Bauer, N., Kiefer, F. 2024. Genetically Encoded Reporters to Monitor Hypoxia. In: Gilkes, D.M. (eds) Hypoxia. Methods in Molecular Biology, vol 2755. Humana, New York, NY.

45. Pulido, R., López, J.I., Nunes-Xavier, C.E. 2024. B7–H3: a robust target for immunotherapy in prostate cancer. Trends Cancer 10(7):584–7.

46. Shen, Q., Zhou, K., Lu, H. et al. 2024. Immune checkpoint B7-H3 is a potential therapeutic target in prostate cancer. Discov Onc 15, 822.

47. Zhao, S., Zhang, H., Shang, G. 2025. Research progress of B7-H3 in malignant tumors. Front. Immunol., 16.

48. Nguyen, P., Okeke, E., Clay, M., Haydar, D., et al. 2020. Route of 41BB/41BBL Co-stimulation Determines Effector Function of B7-H3-CAR.CD28z T Cells. Molecular Therapy: Oncolytics Vol. 18.

49. Moya, L., Walpole, C., Rae, F., et al., 2023. Characterisation of cell lines derived from prostate cancer patients with localised disease. Prostate Cancer Prostatic Dis. 26(3):614–624.

50. Takahashi, K., Tanabe, K., Ohnuki, M., Narita, M., Ichisaka, T., Tomoda, K. and Yamanaka, S. 2007. Induction of Pluripotent Stem Cells from Adult Human Fibroblasts by Defined Factors. Cell 131(5): 861–872.

51. Mahmoudi, S. et al., 2019. Heterogeneity in old fibroblasts is linked to variability in reprogramming and wound healing. Nature 574: 553–558.

52. E. Ascic et al., 2024. In vivo dendritic cell reprogramming for cancer immunotherapy. Science 386.

53. Kongsbak, M., Levring, T.B., Geisler, C., Marina Rode von Essen, M.R. 2013. The vitamin D receptor and T cell function. Front. Immunol., Sec. T Cell Biology 4.

54. Berdiaki, A., Neagu, M., Tzanakakis, P., Spyridaki, I., Pérez, S., Nikitovic, D. 2024. Extracellular Matrix Components and Mechanosensing Pathways in Health and Disease. Biomolecules 14, 1186.

55. Hotamisligil, G.S. and Davis, R.J. 2016. Cell Signaling and Stress Responses. Cold Spring Harb Perspect Biol 8:a006072 Hanahan, D., Weinberg, R. Hallmarks of Cancer: The Next Generation. Cell, 144: 646–674

56. Hanahan, D. & Weinberg, R.A. 2011. Hallmarks of cancer: the next generation. Cell 144, 646–674.

57. Öhlund, D., Elyada, E. & Tuveson, D. 2014. Fibroblast heterogeneity in the cancer wound. J Exp Med 211, 1503–1523.

58. Jain, R.K. 2005. Normalization of tumor vasculature: an emerging concept in antiangiogenic therapy. Science 307, 58–62.

59. Krysko, D.V. et al. 2012. Immunogenic cell death and DAMPs in cancer therapy. Nat Rev Cancer 12, 860–875.

60. Salmon, H. et al. 2015. A network of tumor-resident dendritic cells, macrophages and monocytes restricts T-cell immunity in cancer. Cell 161, 802–816

